# Just a phase? Causal probing reveals spurious phasic dependence of sensory perception

**DOI:** 10.1101/2023.08.21.554096

**Authors:** M. Vinao-Carl, Y. Gal-Shohet, E. Rhodes, J. Li, A. Hampshire, D. Sharp, N. Grossman

## Abstract

For over a decade, electrophysiological studies have reported correlations between sensory perception and the phase of spontaneous pre-stimulus brain oscillations. To date, these findings have been interpreted as evidence that the brain uses neural oscillations to sample and predict upcoming stimuli. Yet, evidence from simulations have shown that analysis artefacts could also lead to spurious pre-stimulus oscillations that appear to predict future brain responses. To address this discrepancy, we conducted an experiment in which visual stimuli were presented timed to specific phases of spontaneous alpha and theta oscillations. This allowed us to causally probe the role of ongoing neural activity in visual processing independent of the stimulus-evoked dynamics. Our findings did not support a causal link between spontaneous alpha / theta rhythms and behaviour. However, spurious correlations between theta phase and behaviour emerged offline using gold-standard acausal time-frequency analyses. These findings demonstrate that care should be taken when inferring causal relationships between neural activity and behaviour using acausal analyses.

**HIGHLIGHTS:**

- We causally probed the role of spontaneous EEG rhythms in visual attention via phase-locking.
- Theta phase predicted behaviour offline, but cues presented in real-time with theta had no effect on behaviour.
- Spurious theta rhythmic sampling is an artefact of the evoked potential and acausal filtering.
- ecHT accurately computes the phase in real-time and mitigates erroneous phase-behaviour correlations.

## INTRODUCTION

Extensive evidence from electrophysiological studies suggests that sensory perception occurs in cycles (1–7). Attentional awareness of visual stimuli fluctuates with the phase of neural activity at stimulus onset (8–15); choice outcomes during perceptual decision making track the underlying parietal delta and alpha rhythms (16,17); and the strength of memory formation waxes and wanes with the phase of spontaneous gamma oscillations across the visual cortex (18). Behavioural periodicities have been reported for a myriad of tasks (19–22), in humans and animal studies (21,23), and across sensory systems (20,24–28).

To date, these findings have been interpreted as evidence that the brain uses neural oscillations to sample and predict upcoming stimuli (12,29,30). However, it has been debated whether these oscillations could also be introduced artefactually during time-frequency (TF) analysis (31–34). For example, applying a filter to a transient brain response can lead to oscillations or ripples around the event known as “ringing”. This occurs due to the oscillatory response of a filter to an impulse-like input. These operations can also smear out the brain response in time. Time-domain shifts can arise when the output of a TF analysis depends on both past and future input values. These operations are termed *acausal* because they violate causality assumptions (there is no natural system where the past is influenced by the future). **Figure 1A-B** illustrates this phenomenon.

**FIGURE 1.**
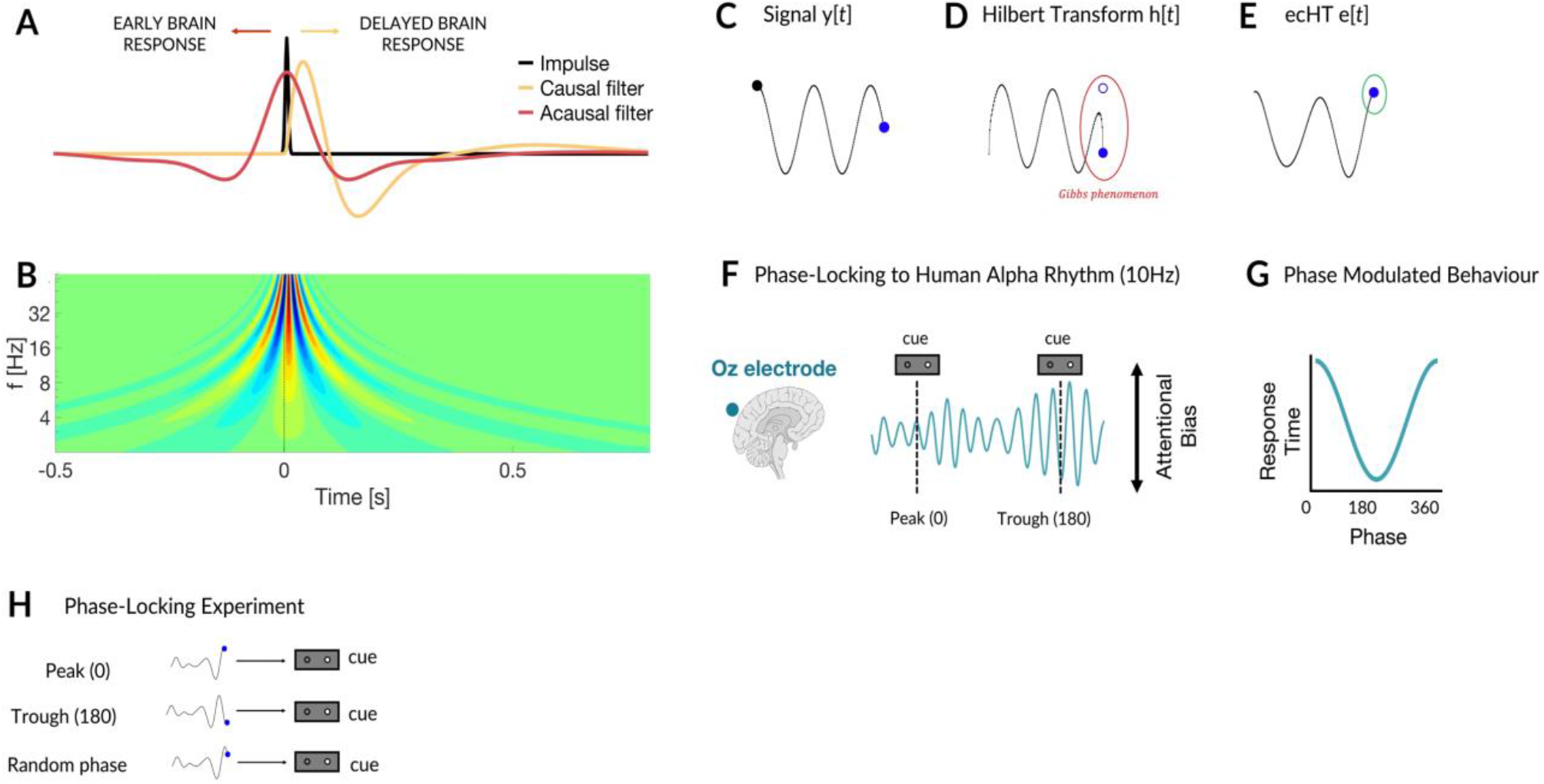
THE PHASE-LOCKING CONCEPT AND TIME-FREQUENCY ARTEFACTS. Brain responses to sensory events are often brief and intermittent in time. In some cases filtering an evoked response can smear it in the time domain so that it appears before the stimulus has even been presented [1.A]. Filtering around an impulse-like event can also evoke spurious oscillations (or “ringing” effects) [1.B]. A ringing artefact which has been smeared backwards in time could present as a pre-stimulus oscillation that predicts future behaviour. **TEMPORAL DISTORTIONS THROUGH TIME-FREQUENCY ANALYSES** A. Delayed response of a causal bandpass filter and early response of a acausal bandpass filter applied to an impulse signal. Both filters 2nd order Butterworth 1-10Hz). B. Ringing effect produced by wavelet analysis of an impulse signal generating spurious oscillations across multiple frequencies before the onset of the impulse. **PHASE-LOCKING CONCEPT** C-D. Computation of the analytic signal using the Hilbert transform leads to distortions in the end points of the signal. E. The end-point corrected Hilbert transform (ecHT) applies a causal bandpass Fourier filter to the DFT of the analytic signal, restoring the oscillatory properties at the end points of its Fourier representation, rending accurate computation of the phase. **RHYTHMIC THEORY OF ATTENTION** F-G. Attenuation of visual processing is often reported at alpha peak and enhanced at alpha trough, leading to fluctuations in response time or hit rates to a visual cue H. In the phase-locked choice reaction task visual, cues on a given trial were presented in one of three randomised conditions; either timed to the peak (0), trough (180) or a random phase of the ongoing EEG signal.

Many acausal operations such as zero-phase filters, Hilbert transform, and wavelet convolution are fundamental for analysing and interpreting time-series data. However, they complicate inferring causal relationships between brain activity and behaviour. For example, a brain response which has been smeared backwards in time could be misinterpreted for a pre-stimulus oscillation that predicts future behaviour.

Simulations have demonstrated that indeed inappropriate TF analysis can lead to smearing of stimulus-evoked activity in the time domain (32), and it has been suggested that such effects could distort relationships between the ongoing phase dynamics and behaviour (31). However, to date empirical testing of a link between ongoing phase dynamics and behaviour has not been reported using causal analyses.

There are two operations which make gold-standard TF analyses acausal. First, the instantaneous phase and amplitude are usually computed using acausal operations such as the Hilbert transform or wavelet convolution. Second, the pre-stimulus TF window overlaps with the evoked brain response. This cannot be avoided because aligning the stimulus onset with the end of the TF analysis window leads to distortions that near the window edges after applying the Hilbert transform (so called Gibbs distortions). This means that the phase at the onset is typically computed with acausal TF operations within a window that overlaps with the stimulus-evoked response.

Here we address this long-lasting debate by directly probing the spontaneous phase dynamics independent of the stimulus-evoked dynamics. To do this we designed an experiment where the visual inputs were presented timed to the phase of ongoing alpha and theta oscillations recorded using electroencephalography (EEG). As discussed previously, phase-locked experiments are challenging because computing the phase in real-time via the Hilbert transform (HT) leads to Gibbs distortions at the end-points of the signal (35). Until recently this meant that the instantaneous phase could not be computed in real-time. To solve this, we developed the end-point corrected Hilbert transform (ecHT) **(Figure 1C-E)**, a simple yet powerful method that tracks the instantaneous phase and envelope amplitude of an oscillatory signal (36). In our phase-locked experiment visual cues were presented at the real-time peak and trough of theta oscillations (condition 1), or alpha oscillations (condition 2), as previous studies have widely reported bidirectional gain on visual perception at these phases (37–40) **(Figure 1F-G)**.

## RESULTS

### Phase locked visual choice reaction task

Single channel EEG was recorded from 20 healthy subjects (mean age = 22.9, std =2.17, F= 13) performing a phase-locked choice reaction task (CRT) Figure.1B. The task was conducted on a custom-made device capable of recording a single channel EEG, tracing the instantaneous phase of the signal in real-time using the ecHT, and synchronising visual cues (i.e., left/right LED arrows) to pre-determined phases of the ongoing EEG signal. This closed-loop experimental design allowed us to test the prediction that response times (RTs) would be modulated by the phase at which the visual cue was presented.

In the experiment, visual cues were either phase-locked to the frontal midline theta-band oscillation (5Hz) recorded from Fz (condition 1), or to the occipital alpha-band oscillation (10Hz) recorded from Oz. (condition 2). Cues were delivered in one of three pseudo-randomised phase-locked conditions — peak (0), trough (180), or a random phase (sham) **(Figure 1H)**. Phase-locking accuracy, as measured by the resultant vector length, was maximal at the target frequencies **(SFigure 1A)** and did not differ between peak and trough conditions **(SFigure 1B)**.

### Behavioural performance under phase locked conditions

We found that actively presenting cues in time with the peak or trough did not lead to a difference in RT, either between phase-locked conditions, or between phase-locked conditions and sham **(Figure 2A)**. We also observed no difference between conditions after modelling the effects of inter-stimulus interval and hysteresis **(see methods)**.

**FIGURE 2.**
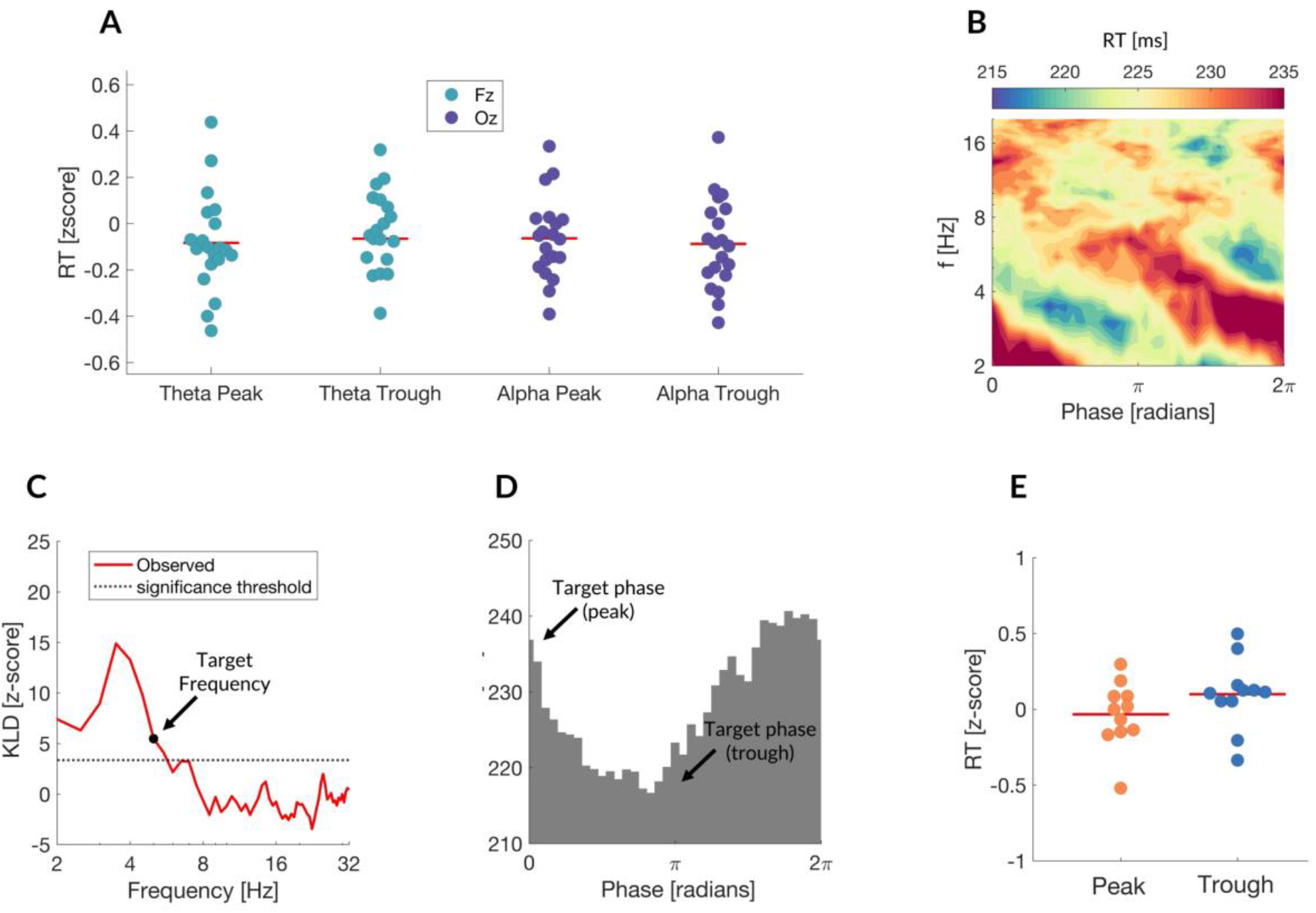
NO CAUSAL RELATIONSHIP BETWEEN RESPONSE TIME AND ONGOING PHASE. **BEHAVIORAL RESULTS** A. We observed no difference in performance either between phase-locked conditions or against sham. Median-normalized group-level RTs (red bar indicates bootstrapped median); green and purple dots depict individual participants from experiment 1 (Theta-Fz) and experiment 2 (Alpha-Oz) respectively. B. phase-resolved performance for single exemplar subject at cue onset. RTs are modulated by low frequency (2-6Hz) phase at single electrode Fz. C. Group level Kullback-Leibler divergence (KLD) for each frequency. KLD is calculated for each frequency and normalised to a surrogate distribution where trials were shuffled (5000 permutations). Note the peak phase-dependent modulation of performance in the theta band ∼4Hz. KLD was computed from 2–32 Hz in 1Hz steps, dotted line indicates significance threshold corrected for the family wise error rate. D. RT as a function of 5Hz phase for experiment 1 (Theta-Fz) in single exemplar subject. Note that peak (0) and trough (180) of 5Hz phase are approximately associated with maximum and minimum RTs, despite no actual difference in performance in (E) E. No change in performance between orthogonal phase conditions. Only subjects in which phase locked trials had a significant KLD at 5Hz and were within 30° of optimal / suboptimal phase bins, were included (n = 11). Normalised Phase distribution for orthogonal phase conditions in preferred frequency for peak and trough trials. Distributions were created by subtracting the mean of phase condition 1 from both phase conditions.

Sustained attention has previously been linked to alpha and theta oscillations. However, the precise frequency and phase which elicit maximum and minimum gain on performance can be specific to an individual. To account for this, we ran a second analysis only on subjects where peak and trough phases which were targeted predicted modulation of behaviour offline (n=11). **Figure 2B-D** shows a representative subject. However, even in these subjects there was no difference in RT when visual cues were actively presented timed to the peak and trough of alpha and theta oscillations **(Figure 2E)**.

### The effect of causal and acausal analyses on phase – behaviour relationship

How could alpha and theta phase correlate with RT when the phase is computed offline, yet have no influence on behaviour when visual cues were phase-locked in real-time to the same brain oscillations?

Previous literature has demonstrated that the power and phase of stimulus-evoked oscillations can predict behavioural outcomes (41–43). It is also known that acausal filters can lead to a smearing of transient waveforms such as the evoked response in the time domain (31,33,44). This is because the outputs of acausal filters are affected by future inputs, while the outputs of causal filters only depend on past and present inputs. We therefore hypothesised that applying an acausal filter would cause stimulus-evoked activity to correlate with behaviour before the stimulus has occurred, thereby causing ongoing activity to appear to predict future responses.

To test this, we compared the phase-behaviour relationship when the pre-stimulus phase was computed using the ecHT (causal), against the phase computed with the HT (acausal). To compute the phase with the ecHT we apply the algorithm to the pre-stimulus window centred at the edge. The algorithm corrects the phase distortion at the end-points, thus the pre-stimulus phase is resolved independent of the stimulus-evoked dynamics (the ecHT never ‘sees’ the post stimulus response). We then regressed the instantaneous phase, and amplitude computed from both approaches against response time **(see methods)**.

We found that the post-stimulus amplitude and phase were predictive of task performance for both approaches **(Figure 3, SFigure 2)**. However, the pre-stimulus phase and amplitude predicted RT up to 300ms prior to stimulus onset for the HT (acausal) **(Figure 3B, SFigure 2B)**, and not for the ecHT (causal) **(Figure 3A, SFigure 2A)**. These results imply that the correlation between spontaneous theta oscillations and performance was produced by an artefactual smearing of stimulus evoked potential.

**FIGURE 3.**
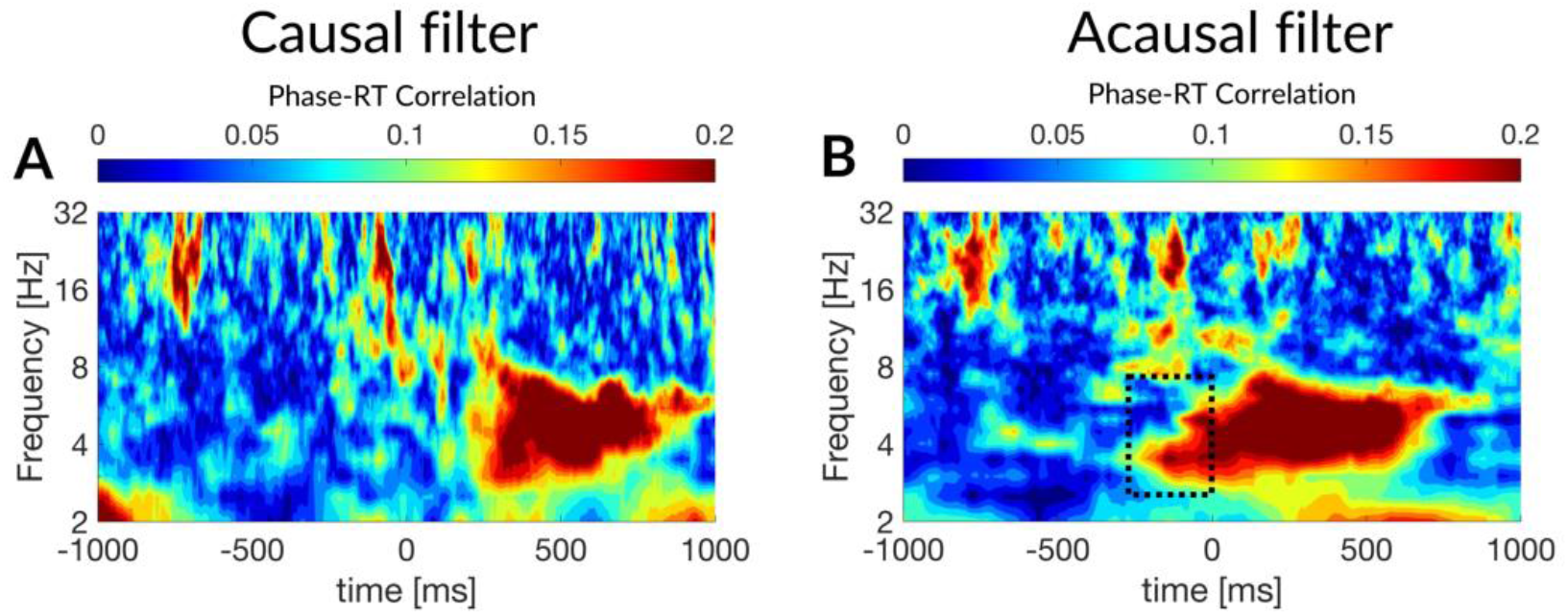
TIME SHIFTED TEMPORAL DYNAMICS OF VISUAL PROCESSING. The temporal dynamics of visual processing are shifted in the time domain depending on the filtering approach. When the phase is computed with the ecHT (causal filter), post stimulus phase predict behaviour (A-B). However, after non-causal filtering and computation of the analytic signal with the Hilbert transform, this relationship is shifted backwards in time to the pre-stimulus. Colorbar indicates coefficient of determination for circular – linear correlation. A. Group level circular-correlation between ecHT-phase and RT. Low frequency phase correlates (3-8Hz) with RT post-stimulus. B. Group level circular-correlation between HT-phase and RT. Phase-RT correlation shifts to pre-stimulus

### Theta phase-dependent behaviour is an artefact of the evoked potential

To directly test whether this artefact is caused by the evoked response, we replaced the pre-stimulus EEG in each trial with white noise, thus destroying any relationship between spontaneous oscillatory activity and behaviour **(Figure 4A)**. However, we still observed a robust relationship between the phase of the white noise signal from 2-8Hz and behaviour when the phase was computed with the HT (acausal) **(Figure 4B,D**). This relationship was not present when the phase was computed using the ecHT (causal) (**Figure 4C-D)**.

**FIGURE 4.**
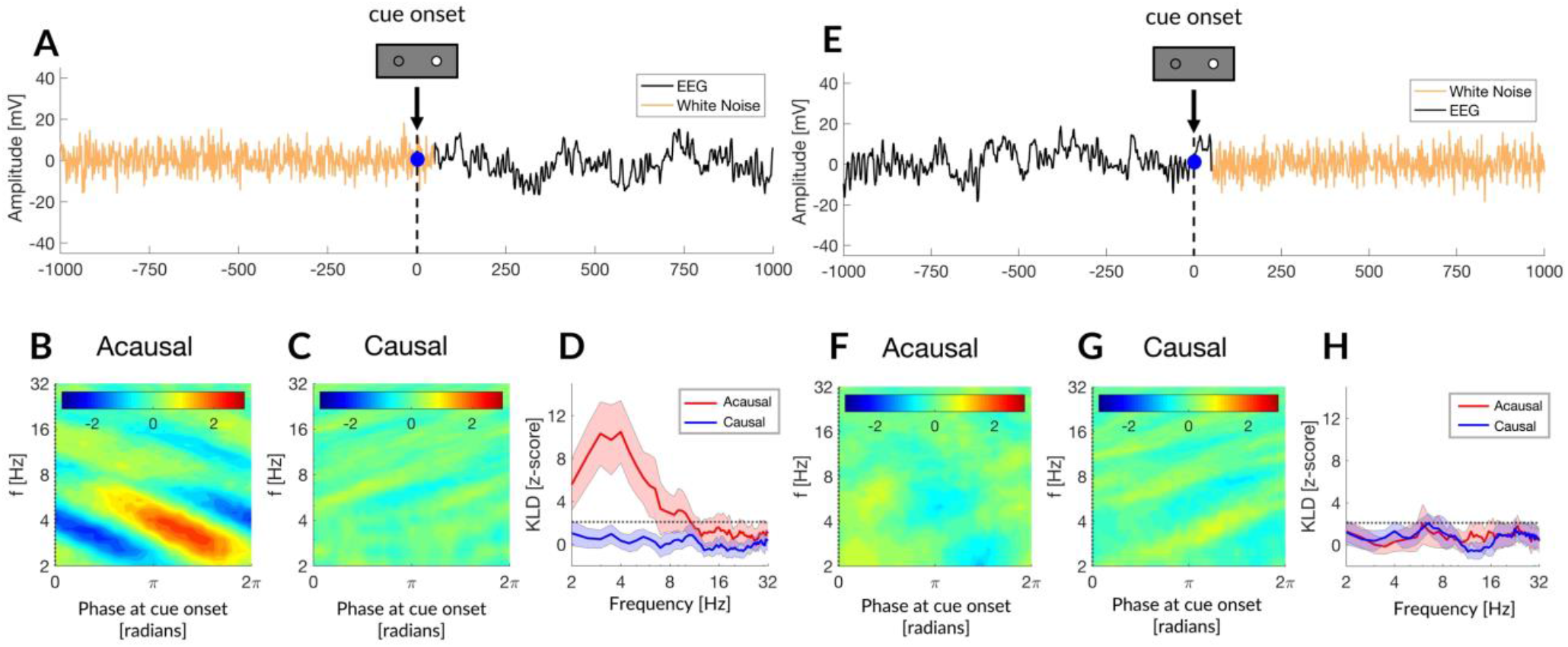
RHYTHMIC SAMPLING AT CUE ONSET IS AN ARTEFACT OF THE EVOKED POTENTIAL. A. Example trial where pre stimulus (−1000:50ms) is replaced with white noise. The Instantaneous phase is computed at cue onset (blue dot) for all trials. B-C. The phase resolved RT was then computed using either the Hilbert Transform (acausal) (B) or ecHT (causal) (C). Note the erroneous relationship between 2-8Hz phase and behaviour when the phase is computed with the Hilbert transform, but not with the ecHT. The ecHT computes the phase using a causal filter which is agnostic to the post stimulus EEG, and correctly identified no relationship between the white noise signal and behaviour (C). D. Normalised KL divergence shown for each frequency band, notice the significance phasic modulation of performance for all low frequencies (2-7Hz) for the Hilbert transform (red), but not the ecHT (ecHT). Dotted line indicates significance threshold. E-H. Same approach as in A-D, only the post-stimulus (50:1000ms) is replaced with white noise. Notice that the relationship between the phase at onset (blue) and behaviour previously uncovered in (B) vanishes once the post stimulus is whitened (F). The causal filtering approach with the ecHT identifies no relationship between the phase of the noise and behaviour in both cases behaviour (C,G).

Next, we applied the reverse analysis and replaced the post-stimulus EEG on each trial with white noise. This destroyed any existing relationship between stimulus-evoked dynamics and behaviour, while leaving spontaneous oscillations present in the pre-stimulus intact **(Figure 4E)**. When we did this, the phase - behaviour relationship uncovered at the onset vanished **(Figure 4F-H)**.

These findings demonstrate that the correlations between spontaneous theta oscillations and behaviour were caused by an artefactual smearing of the behaviourally correlated stimulus-evoked potential.

### Backward smearing of behavioural correlated evoked potential in synthetic data

Next, we used synthetic data to further validate that the evoked brain response can give rise to erroneous correlations between pre-stimulus oscillations and behaviour. We used a synthetic pink noise signal to represent the ongoing EEG and embedded a 5Hz wavelet in the ‘post-stimulus’ to represent an evoked potential. We then introduced a correlation between the latency of the wavelet and a random variable (RV). This allowed us to test if the wavelet would lead to a spurious correlation between the pre-stimulus noise and the RV **(Figure 5A)**.

**FIGURE 5.**
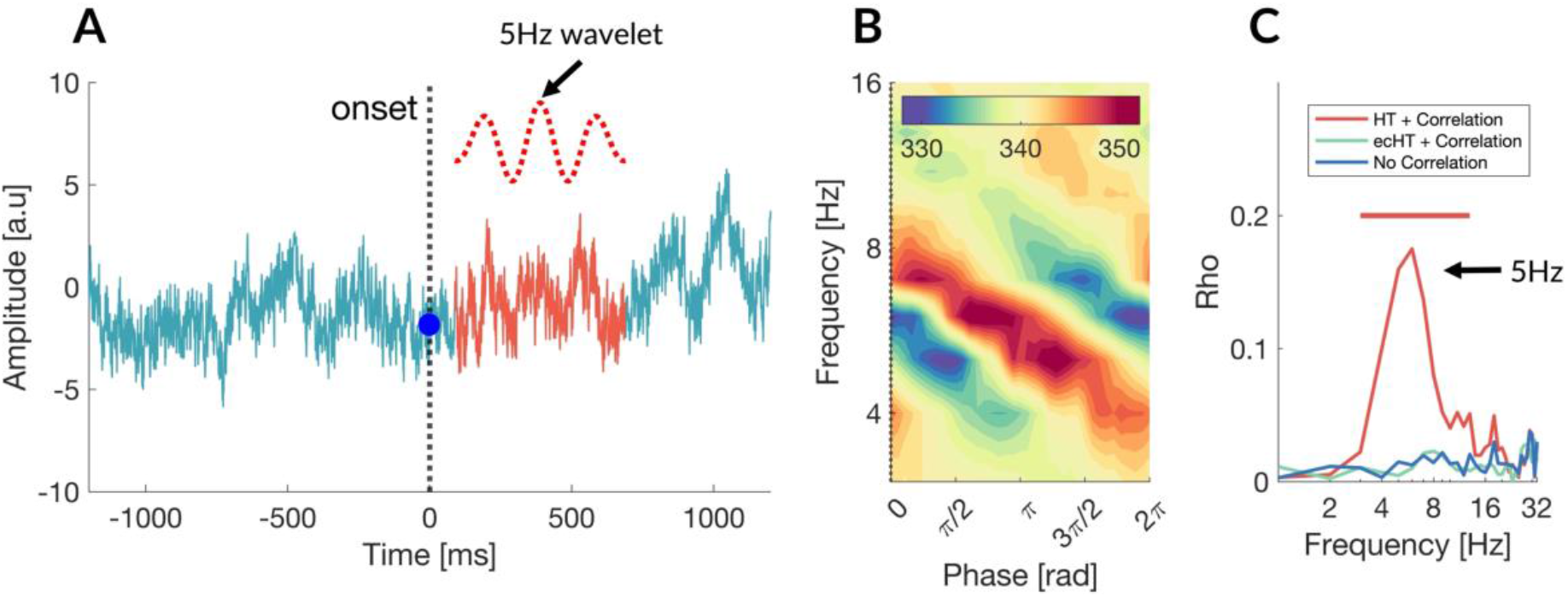
RHYTHMIC SAMPLING AT CUE ONSET IS RELATED TO THE LATENCY OF BRAIN RESPONSES. A. Example pink noise trial with 5Hz Morlet wavelet embedded in the post stimulus at a variable latency (mean 150ms, std = 50ms). The wavelet amplitude was chosen to match the mean amplitude of the pink noise signal at 5Hz. Blue dot indicates the onset (t = 0), where the phase is estimated either using the Hilbert transform or the ecHT. B. Phase-resolved response time at t = 0 computing using the Hilbert transform. Notice the strong rhythmic modulation of RT by the phase of the pink noise at the onset from 4-7Hz. C. Correlation between the phase at onset and RT for all frequencies (2-32Hz) using either the Hilbert transform (red), ecHT (green) or, using Hilbert transform when the correlation between latency and RT is absent (blue). Significance threshold was p < 0.01.

To test the effect of acausal vs causal analysis we computed the phase of the pink noise at the onset of our synthetic trials using either the HT (acausal) or the ecHT (causal). We found that HT smeared the evoked response leading to a spurious relationship between the phase of the noise and the random variable **(Figure 5B-C)**, which was not present with the ecHT or when there was no correlation between latency and the RV **(Figure 5C)**. The acausal HT analysis also led to spurious phasic dependencies in other frequencies (4-11Hz) that were not present in the wavelet, as would be expected due to the ringing effect associated with filters **(Figure 5C)**.

## DISCUSSION

Empirical evidence from electrophysiological studies in humans have shown that sensory perception fluctuates with the phase of pre-stimulus oscillations (1–7). Previously this has been interpreted as evidence that the brain uses neural oscillations to predict upcoming inputs from the environment (12,29,30). In contrast, our findings suggest that some of the evidence supporting this view may erroneously emerge from the way we analyse electrophysiological data. We show that acausal operations implemented during gold-standard TF analyses smear the brain dynamics evoked by the stimulus into the pre-stimuli period. This smearing can lead to an erroneous relationship between behaviour and pre-stimulus brain oscillations.

It is important to note that the strength of this artefact depends on the amplitude / latency of the evoked response and tends to occur with lower frequencies. The artefact also depends on the existence of a true relationship between the evoked brain response and behaviour. Thus, artefactual correlations between the ongoing EEG and behaviour may not be present in experiments where; (1) The evoked response does not predict behaviour; (2) The evoked response is composed mainly of higher frequencies where smearing effects are less pronounced; (3) If there is a long delay in the evoked response such that the pre-stimulus window does not overlap with it.

Our results expand on earlier simulation studies showing the risk of temporal smearing in gold-standard TF analysis practices (31–33,44). The magnitude of the phasic dependency computed in our experiment was ∼15% (i.e., 15% of the variance was explained by the onset phase) consistent with earlier studies reporting phasic dependency of visual processing (5,39).

Although we did not find a significant relationship between the instantaneous phase of spontaneous oscillations and visual performance, future studies exploring different brain regions and perhaps more demanding visual processing paradigms may be able to unveil such relationships. Studies that aim to uncover such relationships using the ecHT must implement the analysis iteratively with the pre-stimulus window aligned so the time point of interest is the last data point. The ecHT is applied to the entire window and the end-point corrected phase is extracted only for the last time-point. The window is then moved forward, and the process repeated for the next time point.

The experimental and computational strategies reported here offer a promising avenue for future investigations into the causal relationship between oscillatory dynamics in the ongoing EEG and behaviour.

## FIGURES

**SUPPLEMENTARY FIGURE 1.**
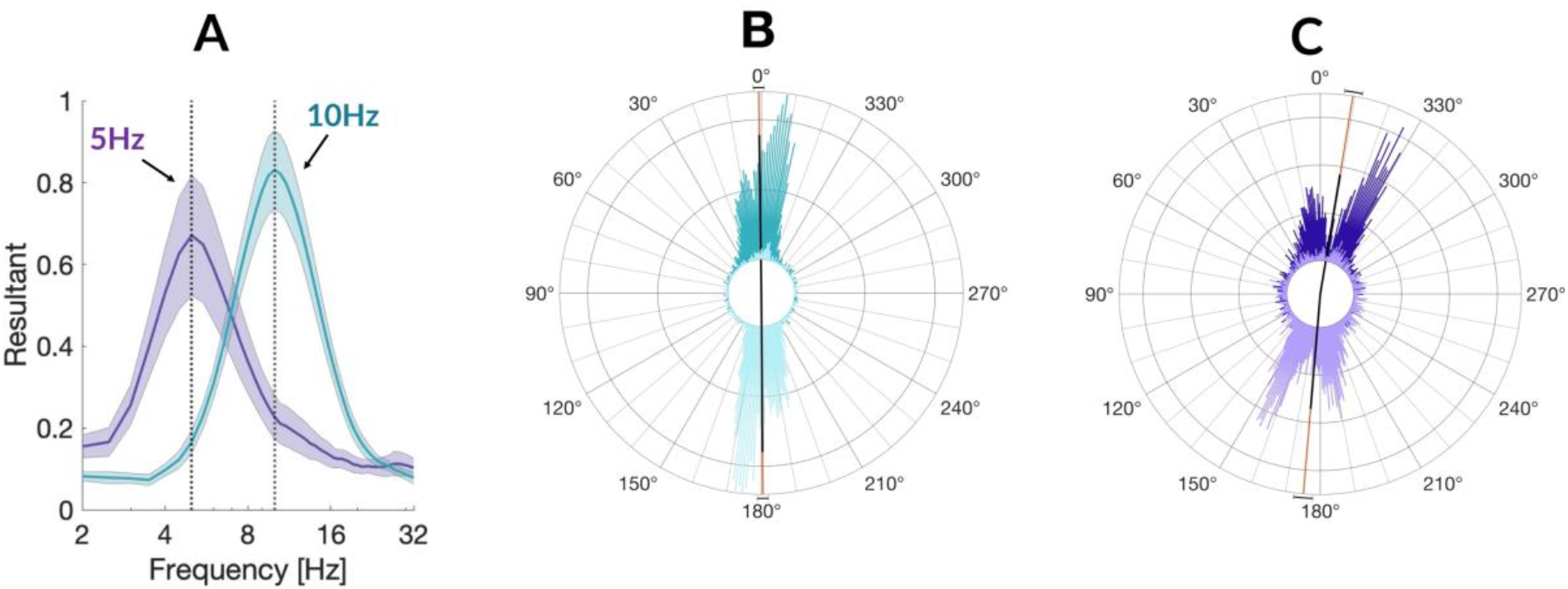
A. Phase locking accuracy to target frequency (5Hz exp1, 10Hz exp2) was quantified by computing the phase resultant vector length at cue onset in the theta band and alpha bands, with a bandwidth of half the central frequency. In both experiments, phase locking accuracy peaked at the target frequency (5Hz, and 10Hz respectively (exp: max phase resultant at 5Hz, mean resultant = 0.66, ci = 0.14; exp2: max phase resultant at 10Hz, mean resultant 0.84, ci = 0.09); Phase resultant in phase –locked conditions differed significantly from sham in both experiments using a paired permutation : Experiment1 : peak v sham t = 7.5, p = 0, trough v sham. T = 14.8, p = 0, 10000 permutations Experiment2 : peak v sham t = 7.3, p = 0, trough v sham. T = 15.3, p = 0, 10000 permutations B. Phase locking accuracy to 0 and 180 degrees for experiment 1 peak mean angle = -9.4 degrees, std = 51.1 degrees. Trough mean angle = 175.2 degrees, std = 52.8 degrees. Phase vector results for peak (shaded dark) and trough (shaded light) conditions also did not differ (p = 0.48, p = 0.64); C. Phase locking accuracy to 0 and 180 degrees for experiment 2 peak mean angle = 0.75 degrees, std = 38.0 degrees. Trough mean angle = 180.0 degrees, std = 37.2 degrees. All phase locked conditions exhibited significant non-uniform phase distributions (ii ‘1; Rayleigh’s test: p = 0, z = 103; iii ‘2; Rayleigh’s test : raleigh’s test p = 0, z = 1.8 103. and different from each other (circular Kuiper test, p = 0, K = 106)

**SUPPLEMENTARY FIGURE 2.**
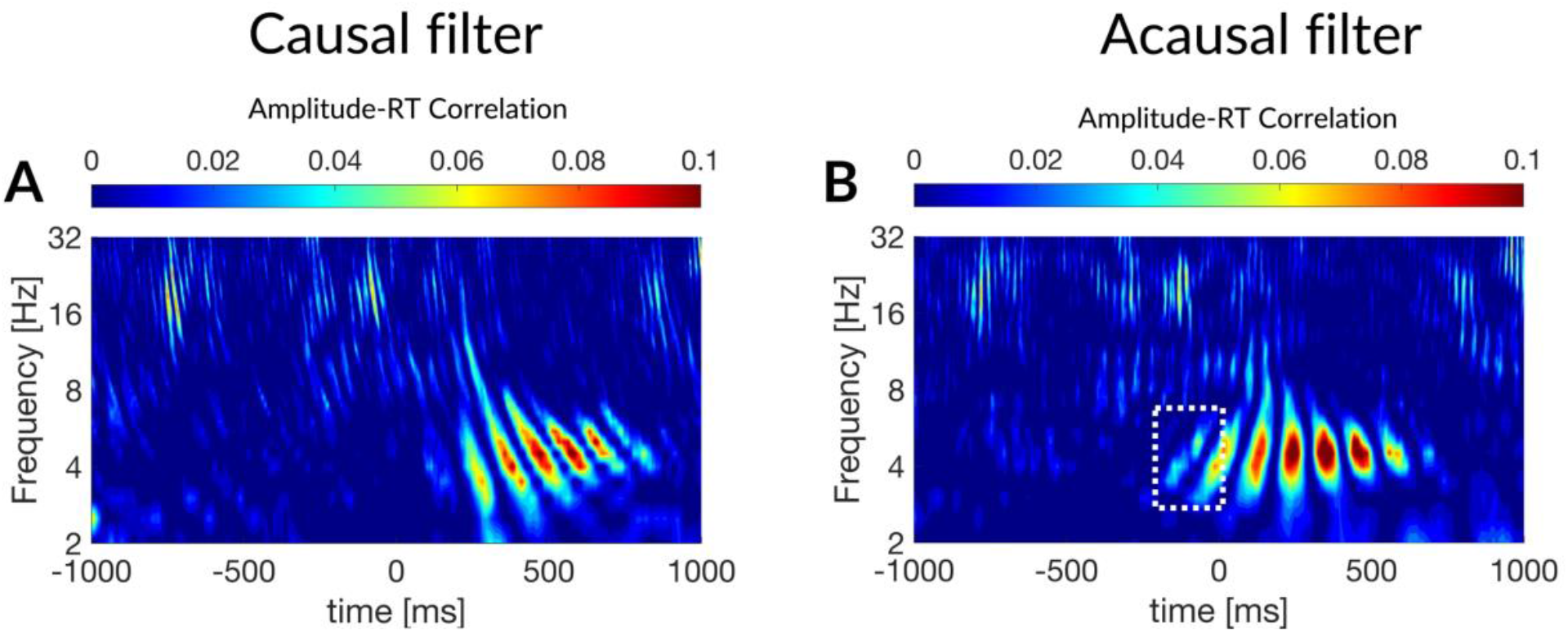
When the amplitude is computed with the ecHT (causal filter), post stimulus amplitude predicts behaviour (A). However, after non-causal filtering and computation of the analytic signal with the Hilbert transform, this relationship is shifted backwards in time to the pre-stimulus (B). Colorbar indicates coefficient of determination for circular – linear correlation. A. Group level regression of ecHT-amplitude against and RT. Low frequency amplitude (3-8Hz) predicts RT post-stimulus. B. Group level regression of HT-amplitude against RT. Amplitude-RT relationship shifts to pre-stimulus.

## METHODS

All analyses were computed in MATLAB.

### Participants, Experimental Design and Procedure

Ethical approval was granted by the Imperial College Research Ethics Committee (ICREC). Participants gave written informed consent and those taking part in the in-vivo experiments were compensated for their time. The study conforms to the Declaration of Helsinki. The experiment was conducted at Imperial College London’s Clinical Imaging Facilities.

Twenty healthy right-handed volunteers (age =22.9 +-2.17, mean + SD=2.17, F= 13) were consenting to participate in a choice response task (CRT). In the CRT, stimulus presentation was controlled with a custom-made response box designed to interface with the closed-loop EEG system. In each trial a visual cue (600 ms) prompting left or right responses was presented on one of two LEDs after variable delay (1600 – 2500ms). Subjects were instructed to give a speeded response to each LED pulse by pressing the appropriate left or right response button. A yellow LED (duration 600ms) in the centre of the response box provided feedback to participants following slow (response times > 600ms from stimulus onset) prompting the participants to increase their response time.

The CRT task was structured into 3 blocks, each bock containing 180 trials and lasting 5m 40s. The total time for running the experiment was 16 minutes including a 30-seconds break between blocks. Each block included 60 trials from each stimulation condition, the order of which was randomised. We continuously recorded EEG dynamics during the behavioural task and delivered each LED pulse according to the randomised stimulation condition, using the ongoing alpha phase decoded from Oz. Randomisation procedures for each block were carried out for (1) the phase-locking condition (sham, peak, trough), (2) for hysteresis of the stimulus type (left or right) and (3) for the inter-stimulus interval (ISI). For sham trials the ISI could take on fixed values from 1600-2100ms in steps of 100ms and was independent of the subject response latency. For stimulation conditions the isi was set to fixed values from 1600-2100ms in steps of 100ms and LED pulse delivered at the earliest target phase after this period had elapsed.

All participants had corrected to normal vision. Exclusion criteria included a history of substance abuse, neurological or psychiatric illness, and a history of or currently prescribed psychoactive medication. The study was approved by the London - West London & GTAC Research Ethics Committee, NHS Health Research Authority.

#### Behavioural Analysis

Performance as measured by response-time: We quantified the performance in each phase-locked condition (peak, trough, and sham) by normalising the RTs on each trial to the median and median absolute deviation within a condition. We then compared the median RT between conditions at the group level by permutation testing. This approach revealing no difference in RT either between phase-locked conditions, or between phase-locked conditions and sham.

Effects of countermanding and ISI: RTs in cognitive control tasks are sensitive to countermanding, also known as sequential effects, and are known to fluctuate with the ISI. We therefore conducted an additional analysis in which ISI and countermanding effects were regressed out using a general linear model. We then compared the median residual variance (RV) between conditions at the group level by permutation testing. We did not observe a difference between phase-locked conditions using either analysis approach, i.e., using RTs or RV as measures of performance.

Preferred frequency and phase: To identify subjects preferred phases we binned the response time by the low frequency phase and computed for each band the pairs of bins with the largest difference in RT. Subjects with mean difference in RT between 0° and 180° phase bins (Z > 1.96) at 5 or 10Hz were defined has having preferred phases at the theta/ alpha peak / trough.

#### End-point corrected Hilbert Transform (ecHT)

Hilbert transform is performed by a discrete Fourier transformation, deletion of the negative frequencies (while compensating to the amplitude loss in the positive frequencies) to get the Fourier representation of the analytic signal (real part, original signal; imaginary part, Hilbert transformed signal), and then inverse Fourier transformation to get the time domain analytic signal.

Yet, the data samples towards the end of the Hilbert transformed signal are contaminated by the Gibbs phenomenon distortion if there is a jump discontinuity between the endpoints of the finite signal. This is since the Fourier transformation of a finite signal represents an infinite repetition of the signal. Thus, a jump discontinuity between the signal ends causes the addition of numerous high frequencies that, when converted to the time domain via inverse Fourier transformation, results in overshooting/undershooting at the ends of the Hilbert transform signal, which leads to erroneous phase estimation.

The ecHT solves this problem by multiplying the discrete Fourier transform of the analytic signal with the frequency response of a causal bandpass filter (a second-order Butterworth filter with a bandwidth of half the central frequency) that effectively smooths the beginning of the signal to the end without distorting the end, thereby removing the jump discontinuity before the inverse Fourier transformation. The ecHT algorithm is fully described in the following paper (36).

#### Causal time-frequency analysis via end-point correction

To compute the phase offline, the ecHT was applied as it would in real-time; by running a sliding window (1s) such that the current time-frequency sample is aligned with the last sample in the window and computing the end-point corrected phase. This allowed us to compute the time-resolved phase independent of the post-stimulus evoked dynamics.

#### Normal time-frequency analysis via Hilbert Transform (acausal)

Acausal computation of the phase and amplitude was conducted by applying a second-order zero phase lag Butterworth filter for a given frequency of interest, with a bandwidth of half the central frequency. The Hilbert transform was then applied to the whole trial and the phase and amplitude extracted from the imaginary and real part of the Hilbert transformed signal respectively.

#### Phase-resolved performance

In order to compute the phase-resolved performance, RTs on each trial were binned by the phase in 90 degree bins in steps of 5 degrees (resulting in 36 RT-phase bins in total), similar to the method used in (37). The RTs were either binned by the ecHT-phase (causal analysis) or by the HT-phase (acausal analysis). The RTs were then permutated against a null distribution in which the RTs were shuffled, to create a normalised distribution phase-resolved performance for each frequency from (2– 32 Hz in 1Hz steps). Permutation testing was conducting using a paired permutation test using the function mult_comp_perm_t1 in MATLAB with 5000, alpha level 0.05.

#### Kullback-Leibler Divergence (KLD)

To test whether RTs were significantly modulated by the phase for a given frequency we computed the KLD for the normalised performance distribution described above, against a uniform distribution. KLD quantifies the amount of information lost when approximating one probability distribution with another with higher values indicating higher dissimilarity between the two distributions. The KLD for each frequency was compared against a null distribution where the RTs were shuffled. Significant frequencies were thresholded at p <0.01.

#### Cluster permutation analysis

To test at what time points phase predict RT we regressed the phase against RT, computed either using the HT (acausal analysis) or the ecHT (causal analysis) for each frequency and time point. This produced a single model for each frequency - time coordinate from 2-64Hz and from 1s pre to 1s post-stimulus. In order to identify significant clusters of time-points and frequencies we performed a cluster permutation analysis. The group level F statistic was compared to a surrogate distribution where the phase and RT were shuffled.

To identify clusters of significant time-frequency points we used the “maximum cluster mass” algorithm, which searches for clusters of contiguous significant points that have the highest total sum of the test statistic within each cluster. Maximum cluster size computed using density based spatial clustering algorithm *dbscan* with the default Euclidean distance metric, epsilon neighbourhood = 2; minimum number of neighbours required for core point = 3.

To Identify significant clusters, we compared the observed F statistic values to the null distribution of test statistics from the permutations. Clusters of contiguous significant points that have a higher total sum of the test statistic than would be expected by chance (p < 0.01).

#### Whitening analysis

For each trial we replaced either the pre-stimulus (−1000:50ms) or post-stimulus (50:1000 ms) with white noise. The amplitude of the white noise signal for each trial was equal to the median amplitude of the raw EEG for that trial. The phase of the hybrid EEG-white noise timeseries was then computed using either the ecHT (causal filter) or the HT (acausal second order Butterworth filter) using the *filtfilt* and *hilbert* functions. The implementation of the ecHT, computation of KLD, cluster correction and permutation analyses are described in the previous sections.

#### Simulation

Correlation between RT and evoked-response latency: To create a weak correlation between RT and latency we used a Gaussian copula to generate correlated pairs of Uniform(0,1) random variables using the MATLAB function *copularnd* (U =copularnd(‘Gaussian’,rho,num_trials), with num_trials = 5000 trials and rho = 0.1 The first column of *U* was extract as RT and the second as the latency. The distribution of RT was transformed to a gamma distribution to match the distribution observed experimentally.

Synthetic EEG: A pink noise EEG signal was generated using the in-built MATLAB function *pinknoise*, and a 5Hz Morlet wavelet was used to capture the impulse-like behaviour of an evoked-response (gaussian width = 7). The wavelet was added to the onset of the simulated EEG signal with a variable latency defined from *U* as described above.

Phase Analysis: The onset phase was computed in 3 ways (1) Hilbert transform on pink noise EEG without the wavelet. (2) Hilbert transform on the wavelet-embedded pink noise signal (3) ecHT on the wavelet-embedded pink noise signal. The implementation was consistent with the analysis used for EEG and behavioral analysis as described throughout the paper.

Circular Correlation Analysis: Similarly, circular correlations between the onset phase and RT was computed using the circular statistics toolbox in the same manner described throughout the paper.

## Contact for Reagent and Resource Sharing

*The data and analysis scripts used in this study will be shared via public repositories before publication*. Further information and requests can be address to the corresponding authors.

## ACKNOWLEDGEMENTS

M.V.C. is funded by the centre for doctoral training (CDT) at Imperial College London through the engineering physical science research council, the UK Dementia Research Institute which is funded by the UK Medical Research Council, Alzheimer’s Society and Alzheimer’s Research UK. N.G. is supported by the UK Dementia Research Institute (UK DRI)—an initiative funded by the Medical Research Council, Alzheimer’s Society and Alzheimer’s Research UK, National Institute for Health and Care Research (NIHR) Imperial Biomedical Research Centre, and the American Alzheimer’s Association.

